# Tillage intensity and plant rhizosphere selection shape bacterial-archaeal assemblage diversity and nitrogen cycling genes

**DOI:** 10.1101/2021.07.16.452714

**Authors:** Mara L. C. Cloutier, Tiffanie Alcaide, Sjoerd W. Duiker, Mary Ann Bruns

**Affiliations:** Ecosystem Science and Management, Pennsylvania State University; Department of Plant Science, Pennsylvania State University

## Abstract

In agriculture, adoption of reduced tillage practices is a widespread adaptation to global change. The cessation of plowing reduces erosion, slows soil organic matter oxidation, and promotes soil carbon accrual, but it can also result in the development of potential N_2_O spots from denitrification activity. In this study, we hypothesized that 16S rRNA-based composition of bacterial-archaeal assemblages would differ in agricultural soils subjected for forty years to a range of disturbance intensities, with annual moldboard plowing (MP) being the most intensive. No-till planting (NT) represented tillage management with the least amount of disturbance, while chisel-disking (CD), a type of conservation tillage, was intermediate. All long-term tillage plots had been planted with the same crops grown in a three-year crop rotation (corn-soybean-small grain+cover crop), and both bulk and rhizosphere soils were analyzed from the corn and soybean years. We also evaluated denitrification gene markers by quantitative PCR at multiple points (three growth stages of corn and soybean). Tillage intensity, soil compartment (bulk or rhizosphere), crop year, growth stage, and interactions all exerted effects on community diversity and composition. Compared to MP and CD, NT soils had lower abundances of denitrification genes, higher abundances of nitrate ammonification genes, and higher abundances of taxa at the family level associated with the inorganic N cycle processes of archaeal nitrification and anammox. Soybean rhizospheres exerted stronger selection on community composition and diversity relative to corn rhizospheres. Interactions between crop year, management, and soil compartment had differential impacts on N gene abundances related to denitrification and nitrate ammonification. Opportunities for managing hot spots or hot moments for N losses from agricultural soils may be discernible through improved understanding of tillage intensity effects, although weather and crop type are also important factors influencing how tillage influences microbial assemblages and N use.

## Introduction

Soil microbial diversity is shaped by abiotic and biotic factors that are constantly changing in agricultural soils subjected to varied management practices. Such practices as nutrient applications, crop rotations, and tillage can impact quality and quantity of soil organic matter (SOM), moisture, temperature, pH, quantity of organic and inorganic nitrogen (N), phosphorous (P), and distribution of crop residues within the soil profile (Elder and Lal, 2008; McDaniel et al., 2014). Management practices can directly or indirectly impact microbial communities by altering nutrient and water supplies, exposing microbes to O_2_ through soil mixing, and changing the types of plant residues introduced into soils (Cookson et al., 2008; González-Chávez et al., 2010; Le Guillou et al., 2019).

Different types of tillage result in varied levels of soil physical disturbance, thus altering microbial habitats and disrupting microbial hyphal connections (Young and Ritz, 2000). Moldboard plowing (MP) inverts soil to a depth of 30-35 cm and requires additional disking and cultivating to prepare soil for planting, thus subjecting soil to the greatest amount of physical disturbance. No-till (NT) involves the least physical disturbance because seeds are introduced into slots in untilled soil while the rest of the soil remains undisturbed. Changing tillage management from MP to NT reduces soil erosion, increases soil water-holding capacity, and slows soil organic matter oxidation to permit more soil carbon accrual, particularly when crop residues are left on the soil surface. Organic matter accumulation on the surfaces of long-term NT soils, however, can increase the risk of soluble nutrient loss in runoff and the development of potential N_2_O spots from denitrification activity (Saha et al., 2021). Other types of conservation tillage practices, such as chisel-disking (CD), mix soils and crop residues to shallower depths of 20-25 cm and represent intermediate disturbance intensities that reduce organic matter oxidation while achieving more uniform nutrient distribution.

Soils directly adjacent to plant roots, deemed rhizosphere soils, induce an additional selection on microbes through root exudation and symbiotic partnerships (see reviews by Bakker et al., 2013 and Pathan et al., 2020). Such variables as organic carbon, N, water content, and O_2_ undergo strong temporal changes in rhizospheres (see review by Hinsinger et al., 2009). Rhizosphere soils also may give rise to important ‘hot-spots’ of activity, which are distinguishable from bulk soils by the increased availability of labile carbon from roots and the competition for nutrients that occurs between microbes and plants (Pathan et al., 2020). Rhizosphere dynamics are not only dependent on the specific crop but can change during crop growth through alterations in root exudations and in the activities of microorganisms in rhizospheres (Li et al., 2014; Houlden et al., 2008; Zhao, et al., 2020a). Addressing how management practices and rhizosphere dynamics synergistically impact microbial communities has important implications for microbial functions and soil and plant health (Schmidt et al., 2019; Zhang et al., 2010; Srour et al., 2020).

In a recent study of management and rhizosphere dynamics of maize, interactions between management (conventional or organic) and soil compartment (rhizosphere or bulk soil) were observed on bacterial community composition, as tracked with 16S rRNA evolutionary gene markers (Schmidt et al., 2019). Bacterial functions tracked based on N cycling genes, on the other hand, were primarily affected by management (Schmidt et al., 2019). Soil bacteria and archaea use N in both assimilatory and dissimilatory processes that affect retention or loss of N from soils (Bhowmik et al., 2017). While denitrification results in loss of N_2_O and N_2_ to the atmosphere (Kuypers et al., 2018), nitrate/nitrite ammonifiers produce ammonium which is retained longer in the soil. As microbial processes may counteract each other, net soil N availability to crops thus depends on relative activities of microbial groups. Gene markers for bacteria that can carry out denitrification (*nirK*, *nirS*), nitrite reduction to ammonium (*nrfA*), and N_2_O reduction to N_2_ (*nosZ*) have all be used in attempts to understand the relationships between soil microbial communities and tillage. These relationships are important for improving agricultural N use efficiency (NUE), or the proportion of soil-available N taken up by crops. Global NUE between 1980 – 2010 was recently estimated to be only about 47% (Lassaletta et al., 2014).

In a meta-analysis of 57 published studies comparing N_2_O emissions and denitrification, it was found that soils from NT treatments had higher numbers of genes for N_2_O production (*nir*) relative to gene numbers for N_2_O conversion to N_2_ (*nos*) when compared to soils from MP tillage (Wang and Zhou, 2020). Authors noted that most of these studies were from short-term tillage experiments. Other studies of long-term tillage practices have found inconsistent results. In some studies, higher N_2_O emissions and greater *nosZ* abundances were observed in soils managed with NT compared to MP (Badagliacca et al., 2018a; Badagliacca et al., 2018b), while others quantified higher N_2_O emissions from tilled soils compared to non-tilled soils (Ussiri et al., 2009; Tellez-Rio et al., 2015a). Differences in results from long-term tillage studies may be due to cropping systems and fertilization treatments (Bayer et al., 2015) or interactions between these factors that affect microbial N use (Liu et al., 2017). Nonetheless, soil in NT treatments have greater water-filled pore space and SOM measured in the top 15-20 cm of the soil profile compared to MP (Grandy et al., 2005; Badagliacca et al., 2018a; Badagliacca et al., 2018b, Ussiri et al., 2009), and these factors have been proposed to be main drivers influencing changes in N cycling dynamics.

Other factors that can influence changes in denitrifier and nitrate ammonifier (*nrfA*) abundances in agricultural soils include the crop grown and the time of sampling, particularly in relation to precipitation events. Changes to exudation profiles during crop development might affect functions of microorganisms in the rhizosphere (Chaparro et al., 2014). Furthermore, rhizospheres of different crops and at various developmental stages support different abundances of denitrifiers and rates of denitrification (Zhao et al., 2020b; Usyskin-Tonne et al., 2020). Effects of crop species and developmental stages on denitrifier and nitrate ammonifier gene abundances have yet to be analyzed in the context of long-term management practices such as tillage.

In the present study, soil bacterial-archaeal community composition and denitrification genes were evaluated in soils from three tillage treatments of varied intensity carried out for 40 years. Objectives of the study were to: (1) Assess how level of disturbance intensity influenced assemblage diversity and relative abundances of N cycling genes associated with denitrification and nitrate ammonification; (2) Investigate how different crop entries, corn and soybean, influenced assemblage diversity (within and across tillage treatments) and relative abundances of N cycling genes in the long-term tillage experiment; and (3) Evaluate how bacterial-archaeal assemblages and denitrification gene abundances changed across different growth stages for corn and soybean. Ratios of functional genes, *nrfA* : *nir* (*nirS*+ *nirK*) and *nir* : *nos* (*nosZI* + *nosZII*) have been shown to significantly correlate with rates of nitrate ammonification, denitrification, and N_2_O emissions (Putz et al., 2018; Wang and Zou, 2020). Soils with higher ratios of *nrfA* : nir have higher rates of nitrate ammonification, while higher ratios of nir : nos correlate with greater production of N_2_O compared to reduction of N_2_O. Therefore, we also included these gene ratios in our analyses.

## Materials and Methods

### Site description and management history

Soil samples for this experiment were collected from plots in a long-term tillage experiment at the Russell E. Larson Agricultural Research Center, Rock Spring, Pennsylvania, USA. A more detailed description of the soils and climate of this site can be found in Duiker and Beegle, (2006). This experiment, first established in 1978, was in continuous corn until 2004 when a 3-year rotation of corn-soybean-small grain/cover crop was implemented. Samples for this experiment were collected in 2018 and 2019 in the corn (variety Pioneer P0506AM) and soybean entries, approximately 40 and 41 years after tillage treatments were established. Corn was planted on May 29, 2018, and soybean was planted on June 12, 2019. Corn was fertilized with an initial amendment of 4 gallons of N-P-K 10-34-0 plus 150 lbs of 21-0-0-24 at planting and side dressed with 40 gallons of 30% UAN on July 3, 2018. Soybeans were inoculated but not fertilized.

Of the tillage practices included in the experiment, we sampled the no-till (NT), chisel-disk (CD), and moldboard plow (MP) treatments to assess effects of varied levels of physical disturbance to the soil profiles. We refer to NT as ‘low’ disturbance intensity, while the CD treatment is referred to as ‘intermediate’ disturbance, where the 20-25 cm tillage depth mixes the upper portion of the soil profile kept in place. The MP treatment is referred to as ‘high’ disturbance intensity because plowing inverts the 30-35 cm layer of soil to bury the top portion. Tillage treatment plots were in a randomized complete block design with four replications. High rainfall in 2019 resulted in flooding of two of the treatments in block 1, therefore, block 1 was not sampled in 2019. Average daily temperatures between June 1 to July 31 in 2018 and 2019 were 20.32°C and 20.76°C, and cumulative rainfall during these periods were 410.97 mm and 149.35 mm, respectively.

### Rhizosphere and bulk soil collection

Samples were collected at three different growth stages of corn and soybean and included corn at vegetative growth stages, V3/4, V5/6, V8/9 corresponding to sampling dates June 14, July 3, and July 13, 2018, and soybean at stages V1, V3, and reproductive stage R1, corresponding to sampling dates July 7, July 23, and July 31, 2019. Rhizosphere samples were collected by excavating plant roots with shovels, removing excess soil from the roots, placing the roots in a Ziploc bag taken to the lab for rhizosphere soil collection. Composite bulk soils were also collected at each of the sampling dates at a depth of 0-15 cm using a soil corer with an inside diameter of 2 cm. Soils were stored on ice and transported to the lab. Soil not tightly adhered to the roots was removed by shaking and the remaining soil was considered rhizosphere soil. Approximately 12 roots and 12 bulk soil samples were collected during each sampling event in each block and mixed to create one homogenized sample per block. Samples were stored at -80°C until further analysis.

A portion of the homogenized bulk soils sampled at the beginning of each year (early May) were air dried and sent to the Penn State Agricultural Analytical Services Laboratory (University Park, PA). Soil fertility and particle size analyses were performed on the samples to assess soil pH (H_2_O), Mehlich III extractable concentrations of P, K, Mg, Ca, Zn, Cu, S, acidity, cation exchange capacity (CEC), CEC-Ca %, CEC-K %, and CEC-Mg %. Values for these measurements are presented in Supplementary Table 1.

### Illumina sequencing

Soil genomic DNA was extracted from 0.25 g of thawed soil rhizosphere and bulk samples using the DNeasy Power Soil Kit (Qiagen, Germantown, MD, USA). Amount and purity of DNA were assessed using a NanoDrop 2000 (Thermo Fisher Scientific, Walthan, MA, USA). A two-step PCR pipeline was used to prepare DNA extracts for 16S rRNA sequencing. First, samples were PCR amplified using the 515F/806R primer pair from the Earth Microbiome Project (https://press.igsb.anl.gov/earthmicrobiome/protocols-and-standards/16s/). Approximately 2 µL of DNA was mixed with 1.5 µL forward primer, 1.5 µL reverse primer, 12 µL 5Prime HotStart MasterMix (Quanta BioSciences Inc., Beverly, MA, USA) and 13 µL of PCR grade water to bring the final volume to 30 µL. Amplification was performed on an Applied Biosystems 2720 Thermo Cycler using the following thermal cycling steps: 95°C for 5 min, 35 cycles of 95°C for 45 s, 55°C for 60 s and 72°C for 60 s, and a final elongation step of 72°C for 5 min and a hold at 4°C.

Amplification of 16S rRNA genes was confirmed by gel electrophoresis, after which amplicons were sent to the Penn State Huck Institutes Genomics Core Facility, where initial PCR products were cleaned and the second PCR step was performed to attach Illumina adaptors. This amplification step included 5 µL of the cleaned PCR product, 5 µL of each forward and reverse Nextera Index Primers, 25 µL KAPA HiFi HotStart ReadyMix, and 10 µL PCR water. Thermal cycling steps for the second PCR were 95°C for 3 min, 8 cycles of 95°C for 30 s, 55°C for 30 s, and 72°C for 30 s, and a final step of 72°C for 5 min. These products were then cleaned, quantified and normalized. Samples were sequenced using an Illumina MiSeq for 250 X 250 bp paired-end sequencing (Illumina, San Diego, CA, USA). Samples were demultiplexed by the facility. A negative control was included in the first PCR step and was included on the MiSeq run.

### Sequence Processing

Amplicon sequences were processed and filtered in R Studio version 1.1.383 using R version 3.6.1 using DADA2 (R Core Development Team, 2020; Callahan et al., 2016). Sequences were filtered and trimmed using ‘truncLen’ set to 230 and 200, ‘maxN’ = 0, ‘maxEE’ set to 2 and 2, and ‘truncQ’ = 2. Primers were also removed at this step by setting ‘trimLeft’ to 20 and 20. Error rates were modeled, samples were dereplicated, and amplicon sequence variants were inferred. Forward and reverse reads were merged and samples less than 250 or greater than 254 bp length were removed. Chimeric sequences were removed and taxonomy was assigned using the silva_nr99_v138 database. Further filtering to remove ASVs not assigned to the Kingdom of Bacteria or Archaea and to remove ASVs assigned to the Family of Mitochondria was performed. Finally, samples with less than 3 000 sequences were removed and included the removal of the negative control.

### Nitrogen cycle marker gene quantification

Extracts of DNA were used for quantitative PCR (qPCR) to quantify *nrfA*, *nirS*, *nirK*, *nosZI*, *nosZII*, and 16S rRNA genes. Primers pairs were as follows: *nrfA*F2aw 5’-CARTGYCAYGTBGARTA-3’ *nrfA*R1 5’-TWNGGCATRTGRCARTC-3’ for *nrfA* (Welsh et al., 2014), *nirS*Cd3a-F 5’-AACGYSAAGGARACSGG-3’ *nirS*3cd-R 5’-GASTTCGGRTGSGTCTTSAYGAA-3’ for *nirS* (Kandeler et al. 2006), *nirK*786cF 5’-ATYGGCGGVCAYGGCGA-3’ *nirK*1040R 5’-GCCTCGATCAGRTTRTGG-3’for *nirK* (Henry et al. 2004, modified by Harter et al. 2014), *nosZI*F 5’-WCSYTGTTCMTCGACAGCCAG-3’ *nosZI*R 5’-ATGTCGATCARCTGVKCRTTYTC-3’ for *nosZI* (Henry et al., 2006), *nosZII*-F 5’CTIGGICCIYTKCAYAC-3’ *nosZII*-R 5’-GCIGARCARAAITCBGTRC-’3 for *nosZII* (Jones et al., 2013), and 341F 5’-CCTACGGGAGGCAGCAG-3’ 534R 5’-ATTACCGCGGCTGCTGGCA-3’ for 16S rRNA genes (Muyzer et al., 1993). Reactions included 2 µL of extracted DNA, 10 µL of FastStart Universal SYBR Green Master (ROX) mix (Roche Diagnostics, Basel, Switzerland), different amounts of primer (see Supplementary Table 2 for specific amounts), and PCR grade water to bring the final volume up to 20 µL. Gene quantifications were performed on an Applied Biosystems 7500 Fast Real-Time PCR system (Foster City, CA, USA).

Standards included genomic DNA for *nrfA* and *nosZI*, linearized plasmids for *nosZII*, *nirS*, and 16S *rRNA*, and gBlocks were used for *nirK* (Hackshaw, 2018). Linearized plasmids were prepared using the TA Cloning Kit Dual Promoter, pCRII with TOP10F’ E. Coli (Cat. No. K2060-01, Invitrogen, Carlsbad, CA, USA). Plasmids were purified using the Monarch PCR & DNA Cleanup Kit (New England BioLabs., Ipswich, MA, USA). Linearized plasmids, genomic DNA, and gBlocks were diluted to 10^8^ in 10 mM TRIS pH 8. Standard curves were created using tenfold serial dilutions of the standards that ranged from either 10^2^-10^7^ or 10^3^-10^8^ copies of the gene templates. Negative controls were included on each plate and samples and standards were run in triplicate. Additional details for qPCR and sources of DNA standards can be found in Supplementary Table 2.

### Statistical analyses

All statistical analyses were performed in R Studio (R Core Team, 2020). A phyloseq object was built for the amplicon sequences and used for downstream analyses (McMurdie and Holmes, 2019). Samples that had fewer than 2000 sequences were removed, which resulted in the removal of two samples out of 128. Samples were aggregated at different taxonomic levels using ‘taxa level’ from the microbiomeSeq package (Ssekagiri, et al., 2017).

Multivariate analyses were performed on the 16S rRNA amplicon sequences at the ASV-level using Bray-Curtis dissimilarity and a Permutational multivariate analysis of variances (PERMANOVA) with the vegan package (McMurdie and Holmes, 2019; Oksanen et al., 2019). Post-hoc pairwise PERMANOVAs were performed using 999 permutations with ‘fdr’ p-value adjustments with the RVAideMemoire package (Hervé, 2020). Community composition was further assessed using principal coordinate analyses (PCoA) and differential abundance analysis was performed at the family-level using corncob in R (Martin et al., 2020).

Soil fertility data were analyzed using a linear mixed effect model with tillage and year as fixed effects and block as a random effect. Linear mixed effect, repeated measures analyses were performed on the 16S rRNA amplicon sequences for alpha-diversity and qPCR gene abundances. Samples for alpha-diversity were rarefied to a depth of 4000 sequences per sample for alpha-diversity analyses. Alpha-diversity was estimated using the phyloseq and microbiome packages to include Shannon Diversity, Richness, and Evenness (McMurdie and Holmes, 2019; Lahti et al., 2017). Linear modeling was performed using the lme4, lmerTest, and car packages and post-hoc analyses were performed using the emmeans packages to include ‘fdr’ p-value adjustments (Bates et al., 2015; Kuznetsova et al., 2017; Fox and Weisberg, 2019; Lenth, 2020). Dependent variables were transformed when necessary, to meet modeling assumptions using the bestNormalize package in R (Peterson, 2019). Additional modeling parameters for univariate tests included adding Block as a random variable and plot as the subject of the repeated measures.

### Conditional inference tree analysis

Abundances of nitrogen cycling genes were further assessed to identify constraining variables and interactions between fixed effects of crop year, tillage treatment, and soil compartment with environmental variables. Environmental variables included in this analysis included precipitation 1, 2, and 3 days prior to sampling, mean daily air temperature, daylight hours, growing degree days (based on 50°C for both corn and soybean), and soil moisture. Conditional inference trees use a regression approach to identify predictor variables with the strongest influence on the response variables (N gene abundances). If a predictor variable was significantly correlated with the response variables, the algorithm would split the predictor variable into two groups. At each stage of the analysis a global null hypothesis of independence between the response variable and the predictor variables was tested and if the hypothesis could not be rejected at a set p-value then the estimation would stop. Analyses were performed using ‘ctree’ from the partykit package in R with p <0.10 and 999 permutations (Hothorn and Zeileis, 2015). Overall performance of the tree was assessed using the caret package (Hothorn et al., 2006; Kuhn, 2020). This approach has been used to identify variables constraining variability in organic matter/carbon, N_2_O, soil inorganic N, and microbial diversity in agricultural soils and crops (Saha et al., 2017; van Wesemael et al., 2019; Finney et al., 2015; Ottesen et al., 2016).

Additional R packages that were used to carry out these analyses include devtools, ggplot2, and BiocManager (Morgan, 2019; Wickham, 2016; Wickham et al., 2020). To better understand measurements associated with microbial ecology and multivariate statistics (i.e alpha-diversity and beta-diversity) see review by Hugerth and Andersson, (2017). Amplicon sequences were deposited in NCBI under the BioProject PRJNA690554. Data and code for this project are available at https://github.com/maracashay/Tillage-16SrRNA-MiSeq.

## Results

### Diversity of bacterial-archaeal assemblages by tillage intensity and crop

Tillage intensity, crop type, soil compartment and the interactions among these variables had significant effects on the compositions of bacterial-archaeal assemblages (Table 1). Beta diversity was more affected by tillage intensity (R^2^ = 0.07) than by crop or compartment (both R^2^ = 0.04) and was also influenced by the interaction between all three variables (R^2^ = 0.03). Community composition differed between bulk and rhizosphere soils for nearly all tillage and crop combinations except for the NT treatment in corn (Figure 1, A-B). Community composition in bulk soils under corn differed across the three tillage treatments, while no differences with tillage were observed when comparing bulk soils under soybean (Figure 1, A-B). Community compositions in rhizospheres of corn and soybean also differed from each other regardless of tillage treatment. Communities in rhizosphere soils from the CD treatment also differed from the other corn rhizosphere communities under NT and MP treatments (Figure 1, A-B).

**Figure 1.**
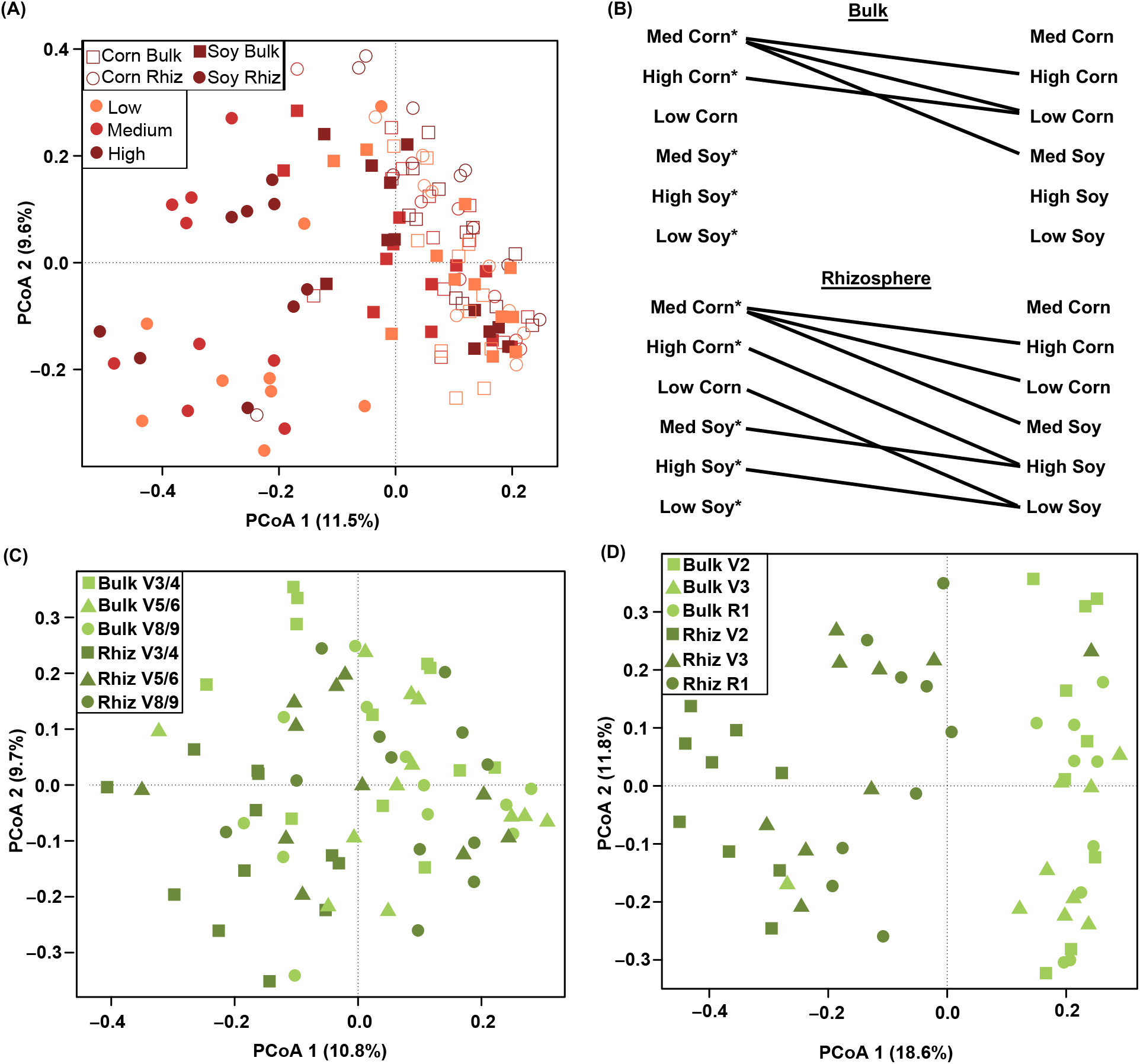
Beta-diversity of the bacterial-archaeal assemblages in the corn and soybean years (A-B), corn only (C), and soybean only (D). Differences in beta-diversity across the two years grouped by crop x tillage x soil compartment are presented in B. Lines connecting sample groups in panel B indicate that those two sample groups have significantly different assemblages. Asterisks on the sample groups on the left-hand side of panel B indicate a significant difference between the bulk and rhizosphere samples for that group.

**Table 1.** Bacterial-archaeal alpha-diversity and assemblage composition (beta-diversity) results from univariate (alpha-diversity) and multivariate (beta-diversity) analyses using data from both corn and soybean years to assess the impact of crop identity, tillage, and soil compartment on community diversity. Beta-diversity was assessed with PERMANOVA and alpha-diversity was assessed using linear mixed effect models. Values presented are the R^2^ and p-values for PERMANOVAs and F-test statistics and p-values from linear modeling and significant values (p <0.05) are marked with an asterisk (*).

| Diversity metric | Alpha-diversity |  |  | Beta-Diversity |
| --- | --- | --- | --- | --- |
|  | Richness | Shannon Diversity | Evenness | Bray-Curtis |
| **Crop** | 47.98, <0.01* | 50.28, <0.01* | 27.53, <0.01* | 0.04, <0.01* |
| **Tillage** | 0.13, 0.87 | 0.63, 0.50 | 0.46, 0.63 | 0.07, <0.01* |
| **Compartment** | 8.57, <0.01* | 22.06, <0.01* | 13.73, <0.01* | 0.04, <0.01* |
| **Crop x Tillage** | 3.09, 0.03* | 5.05, 0.01* | 2.80, 0.06 | 0.02, 0.16 |
| **Crop x Compartment** | 15.28, <0.01* | 20.71, <0.01* | 10.83, <0.01* | 0.04, <0.01* |
| **Tillage x Compartment** | 0.65, 0.52 | 2.28, 0.11 | 0.32, 0.72 | 0.01, 0.22 |
| **Crop x Tillage x Compartment** | 0.10, 0.91 | 1.20, 0.31 | 0.16, 0.86 | 0.03, 0.03* |

Overall, no differences across tillage intensities were observed for alpha-diversity measurements (richness, Shannon diversity, evenness). In contrast, alpha-diversity measures differed for crop type, soil compartment, and interactions between crop type *x* tillage and between crop type *x* soil compartment (Table 1). Alpha-diversity measures were all lower in soybean rhizospheres compared to soybean bulk soils across all tillage treatments, while alpha-diversity in corn rhizospheres and bulk soils did not differ (Supplementary Figure 1). Moreover, community assemblages in soybean rhizospheres had lower richness and Shannon diversity than those in corn rhizospheres across all tillage treatments (Supplementary Figure 1). Across all tillage intensities, corn had higher richness and Shannon diversity compared to soybean (Supplementary Figure 1).

More than 50% of 16S rRNA sequences could be assigned to the ten most abundant families (Figure 2A). Of these ten, five families were also identified as being differentially abundant in bulk soils between tillage treatments. Those families included Chthoniobacteraceae, Nitrosomonadaceae, Pedosphaeraceae, Pyrinomonadaceae, and Sphingomonadaceae. In total, 19 families in bulk soils were differentially abundant across the three tillage treatments. Comparisons between NT and MP treatments resulted in 16 differentially abundant families; NT and CD treatments had 9 families with different abundances, while only three families differed between the CD and MP treatments (Figure 2).

**Figure 2.**
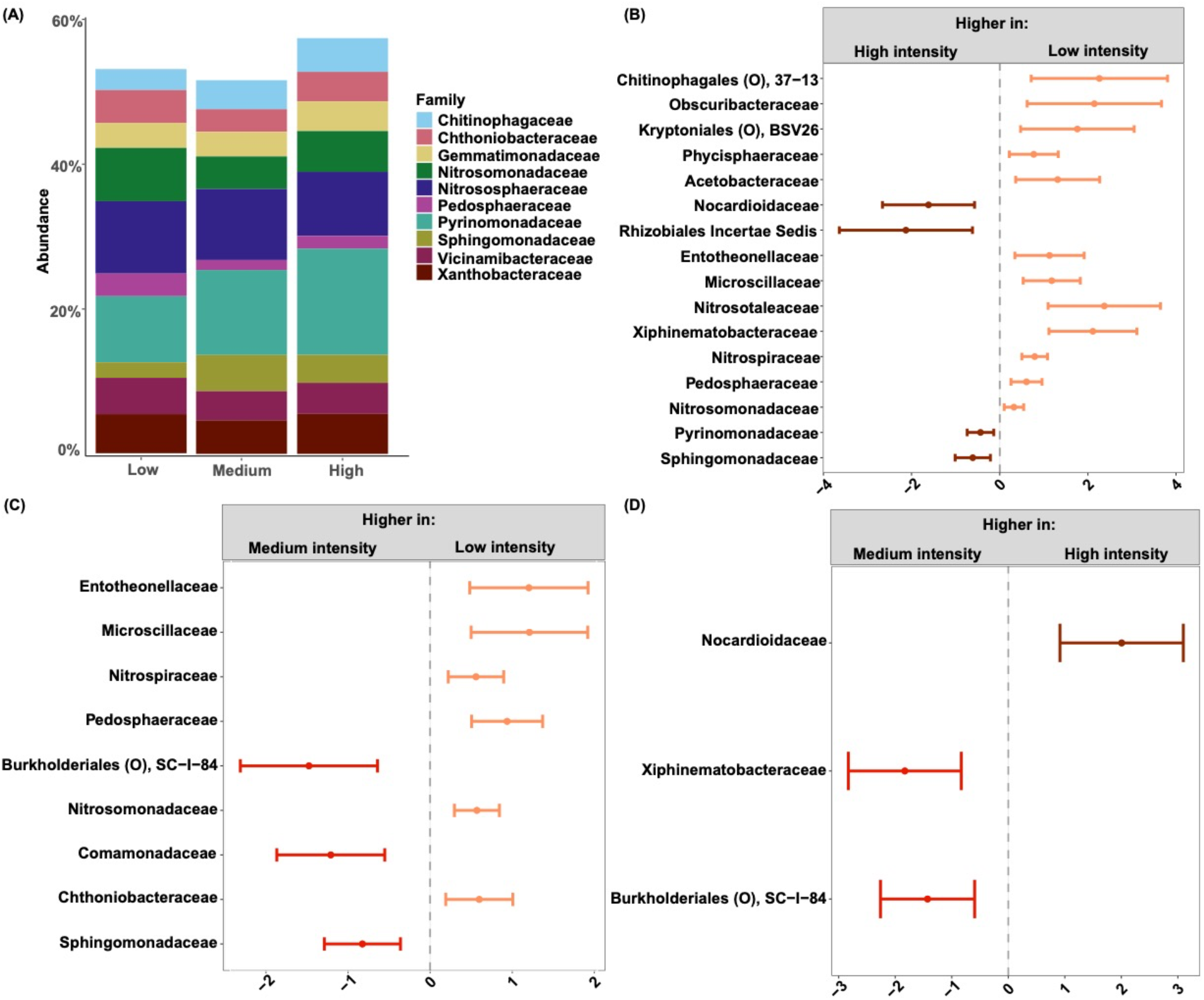
Bulk soil family-level distribution (A) and differentially abundant families taken from both the corn and soybean entries. Families that have differences in abundances between high and low (B), medium and low (C), and medium and high (D) tillage intensities. Values on the x-axis are the coefficient estimates.

### Bacterial-archaeal assemblage diversity across corn growth stages

Beta-diversity of the community assemblages in corn differed by growth stage, tillage, soil compartment, and the interactions between stage *x* soil compartment (Table 1). Of these variables, tillage was the most influential (R^2^ = 0.10). Each growth stage in corn for all tillage treatments had a unique community composition (pairwise PERMANOVA, adjusted p values < 0.05). Assessment of community composition by growth stage *x* soil compartment indicated that bulk and rhizosphere soils from V3/4 differed from the pairs of soils sampled at the V5/6 and V8/9 stages (adjusted p values < 0.05, Figure 1, C). Corn rhizosphere and bulk soils had different community compositions only at stages V3/4 and V8/9 (adjusted p values < 0.05).

Alpha-diversity measurements in corn differed by growth stage, tillage intensity, and the interaction between growth stage *x* tillage (Table 2). Evenness increased with later growth stages, as did richness and Shannon diversity, although the effects were not consistent across tillage treatments (Supplementary Figure 2). Soils managed with MP tillage showed decreased richness at the earliest compared to the latest corn growth stage (V3/4 versus V8/9). In contrast, soils managed with CD tillage showed the opposite trend with increasing richness from V3/4 to V8/9 (Supplementary Figure 2). Richness and Shannon diversity at V8/9 was greatest in the CD tillage treatment compared to either the NT or MP treatments.

**Table 2.** Assemblage composition (beta-diversity) and univariate (alpha-diversity) results from PERMANOVAs and linear mixed effects models using either 2018 (corn) or 2019 (soybean) data to assess the impact of growth stage on community diversity. Values presented are the R^2^ and p-values for PERMANOVAs and F-test statistics and p-values from the linear modeling for alpha-diversity. Significant p-values (p <0.05) are marked with an asterisk (*).

|  | Diversity metric | Beta-diversity | Alpha-diversity |  |  |
| --- | --- | --- | --- | --- | --- |
|  |  | Bray-Curtis ASV-level | Richness | Shannon Diversity | Evenness |
| Corn | Stage | 0.06, 0.01* | 0.20, 0.82 | 0.33, 0.72 | 4.86, 0.01* |
|  | Tillage | 0.10, <0.01* | 2.44, 0.17 | 3.65, 0.03* | 1.91, 0.16 |
|  | Compartment | 0.03, <0.01* | 1.08, 0.30 | 0.15, 0.70 | 0.09, 0.77 |
|  | Stage x Tillage | 0.05, 0.41 | 4.52, <0.01* | 3.97, 0.01* | 1.30, 0.28 |
|  | Stage x Compartment | 0.06, <0.01* | 0.57, 0.57 | 0.50, 0.61 | 0.27, 0.77 |
|  | Tillage x Compartment | 0.02, 0.33 | 0.30, 0.74 | 0.45, 0.64 | 0.11, 0.90 |
|  | Stage x Tillage x Compartment | 0.06, 0.06 | 2.49, 0.06 | 1.97, 0.11 | 0.84, 0.50 |
| Soybean | Stage | 0.05, 0.01* | 0.80, 0.46 | 2.66, 0.08 | 2.23, 0.12 |
|  | Tillage | 0.08, <0.01* | 1.56, 0.22 | 1.81, 0.18 | 1.44, 0.25 |
|  | Compartment | 0.14, <0.01* | 22.62, <0.01* | 33.25, <0.01* | 31.44, <0.01* |
|  | Stage x Tillage | 0.05, 0.89 | 0.98, 0.43 | 0.96, 0.44 | 0.52, 0.72 |
|  | Stage x Compartment | 0.06, <0.01* | 0.96, 0.39 | 1.32, 0.28 | 5.89, 0.01* |
|  | Tillage x Compartment | 0.05, 0.03* | 0.82, 0.45 | 1.06, 0.36 | 0.67, 0.52 |
|  | Stage x Tillage x Compartment | 0.05, 0.78 | 1.41, 0.25 | 1.17, 0.34 | 0.63, 0.64 |

### Bacterial-archaeal assemblage diversity across soybean growth stages

Beta-diversity of bacterial-archaeal assemblages from soybean differed by growth stage, tillage intensity, soil compartment and the interactions among stage *x* soil compartment and tillage *x* soil compartment (Table 2). Of these variables, soil compartment had the greatest influence on assemblage composition (R^2^ =0.14). With respect to the interaction between growth stage *x* soil compartment, differences in assemblage composition were observed between bulk and rhizosphere soils at all growth stages (pairwise PERMANOVA, adjusted p values < 0.05; Figure 1, D). No differences were observed in soybean bulk soils across growth stages (adjusted p values > 0.05). Soybean rhizospheres at stage R1 had a different composition compared to both the rhizospheres at V2 and V3 stages (adjusted p values < 0.05).

Alpha-diversity in soybean soils differed by soil compartment, with evenness affected by the interaction between soil compartment *x* growth stage (Table 2). Both richness and Shannon diversity were lower in soybean rhizospheres compared to bulk soils across all tillage intensities. Evenness was significantly lowest in the soybean rhizospheres at growth stage R1 compared to V2 rhizospheres or R1 bulk soils (Supplementary Figure 2).

### Denitrifier and nitrate ammonifier gene abundances by crop and tillage intensity

Across two crop years, three tillage intensity treatments, and two soil compartments, abundances of denitrifier and nitrate ammonifier genes differed by several orders of magnitude, in descending order: *nosZII* > *nirK* > *nosZI* > *nirS* > *nrfA* (Table 3). Mean ratios of *nir* (NO_2_^-^ to NO) to *nos* (N_2_O to N_2_) were < 1, which may indicate potential for more complete denitrification across these soils (Table 3). Abundances of nitrate ammonification genes (*nrfA*) were much lower than abundances of *nir* genes, with mean ratios < 0.05 in all samples. Gene abundances differed by crop year, tillage treatment, soil compartment and the respective interactions among these variables (Table 3 and Figure 3). Both *nirS* and *nirK* gene abundances were higher in the soybean than in the corn year, while *nosZI* and *nosZII* genes were more abundant in rhizospheres compared to bulk soils overall. The ratios of *nrfA* : *nir* genes were highest in corn rhizospheres and the NT tillage treatments (Figure 3).

**Figure 3.**
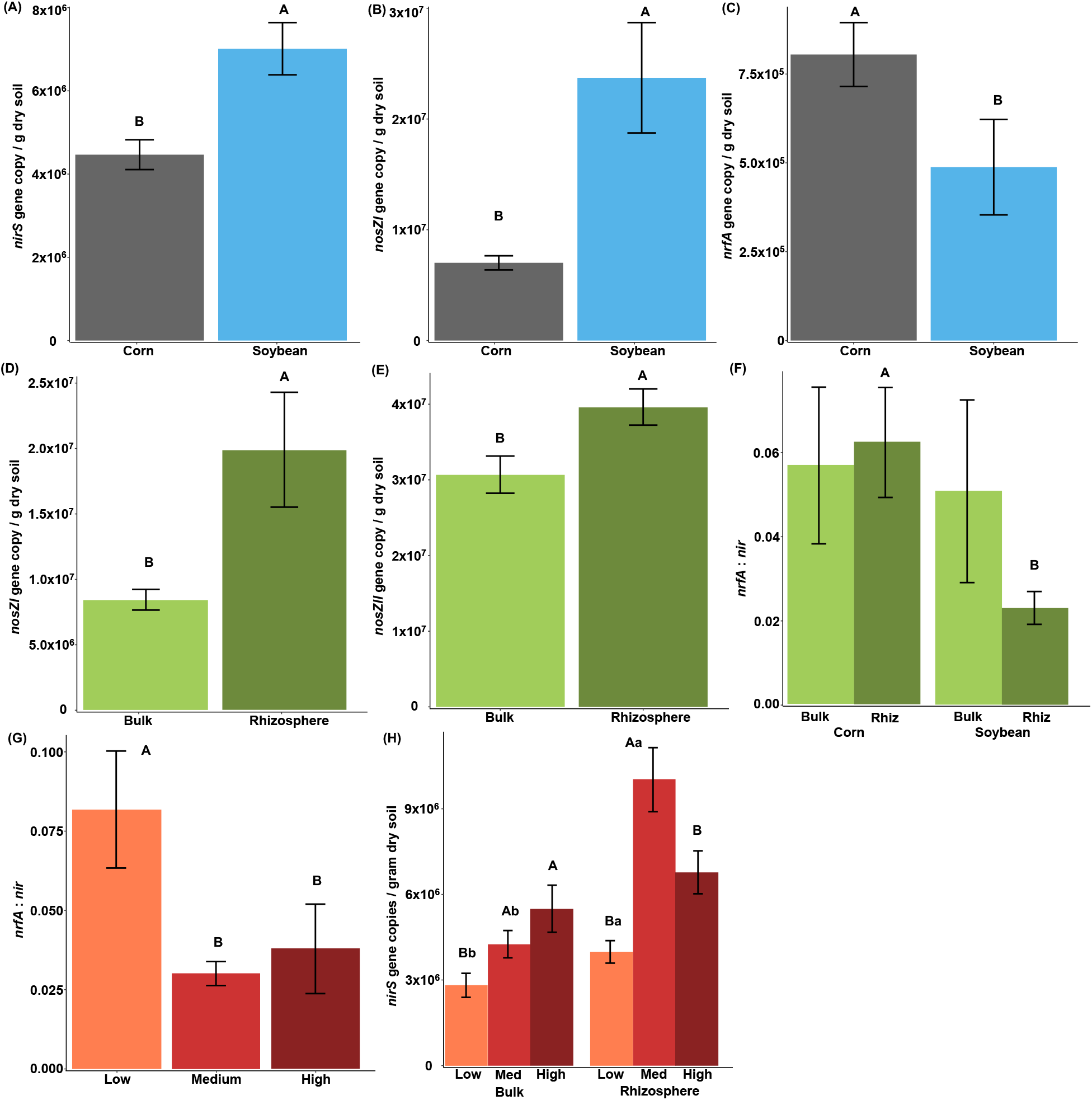
Abundances of genes representing denitrifiers and nitrate ammonifiers from corn and soybean years. Abundances of nirS (A), nosZI (B), and nrfA (C) between crop years. Abundances of nosZI (D), and nosZII (E) by soil compartment and nrfA : nir by soil compartment x crop (F). Abundances of nrfA : nir by tillage intensity (G) and nirS (H) by tillage x soil compartment. Uppercase letters in A-G indicate differences between the treatments. Uppercase letters in H indicate differences between tillage intensity treatments within soil compartment and lowercase letters indicate differences between soil compartment within tillage intensity treatments. Error bars represent standard errors.

**Table 3.** Results of linear mixed effect models for individual N gene abundances. Values presented are the F-and p-values for the linear models and significant variables are denoted with an asterisk (*). Mean average abundances of the genes and gene ratios are also presented.

| Variable(s) | *nirS* | *nirK* | *nosZI* | *nosZII* | *nrfA* | *nir:nos* | *nrfA:nir* |
| --- | --- | --- | --- | --- | --- | --- | --- |
| **Mean averages** | 5.56 x 10<sup>6</sup> | 1.92 x 10<sup>7</sup> | 1.42 x 10<sup>7</sup> | 3.52 x 10<sup>7</sup> | 6.69 x 10<sup>5</sup> | 0.74 | 0.05 |
| **Crop** | 25.26, <0.01* | 0.97, 0.33 | 50.84, <0.01* | 1.58, 0.21 | 15.60, <0.01* | 5.27,<br>0.02* | 5.10, 0.03* |
| **Tillage** | 19.71, <0.01* | 0.09, 0.91 | 0.93, 0.40 | 1.36, 0.26 | 2.70, 0.07 | 1.84, 0.16 | 7.10, <0.01* |
| **Compartment** | 32.56, <0.01* | 1.09, 0.30 | 20.82, <0.01* | 7.55, <0.01* | 2.95, 0.09 | 0.00, 0.97 | 0.12, 0.73 |
| **Crop x Tillage** | 1.87, 0.16 | 0.67, 0.51 | 0.30, 0.75 | 1.12, 0.33 | 0.84, 0.43 | 0.10, 0.90 | 1.56, 0.21 |
| **Crop x**<br>**Compartment** | 0.79, 0.37 | 1.65, 0.20 | 3.07, 0.08 | 2.87, 0.09 | 1.06, 0.30 | 0.45, 0.50 | 4.63, 0.03* |
| **Tillage x**<br>**Compartment** | 4.20, 0.02* | 0.01, 0.99 | 1.85, 0.16 | 1.54, 0.22 | 0.61, 0.54 | 0.29, 0.75 | 0.04, 0.97 |
| **Crop x Tillage x**<br>**Compartment** | 1.26, 0.29 | 0.19, 0.83 | 0.95, 0.39 | 1.22, 0.30 | 1.80, 0.17 | 0.24, 0.79 | 0.74, 0.48 |

### Denitrification gene abundances across corn growth stages

Denitrifier and nitrate ammonifier gene abundances differed by growth stage, tillage intensity, soil compartment, and the respective interactions among these variables (Table 4). Higher abundances of *nirK* and *nrfA* and higher ratios of *nir* : *nos* were observed during the earlier stages of corn growth, while ratios of *nrfA* : *nir* continued to increase over the growing season (Figure 4). Soils managed with NT treatment exhibited higher abundances of *nrfA* compared to MP tillage, and ratios of *nrfA* : *nir* were highest in the NT treatment (Figure 4). Abundances of *nirS* and *nosZI/II* were highest in the CD tillage treatment in corn rhizospheres and were generally lower in the NT treatment (Figure 4).

**Figure 4.**
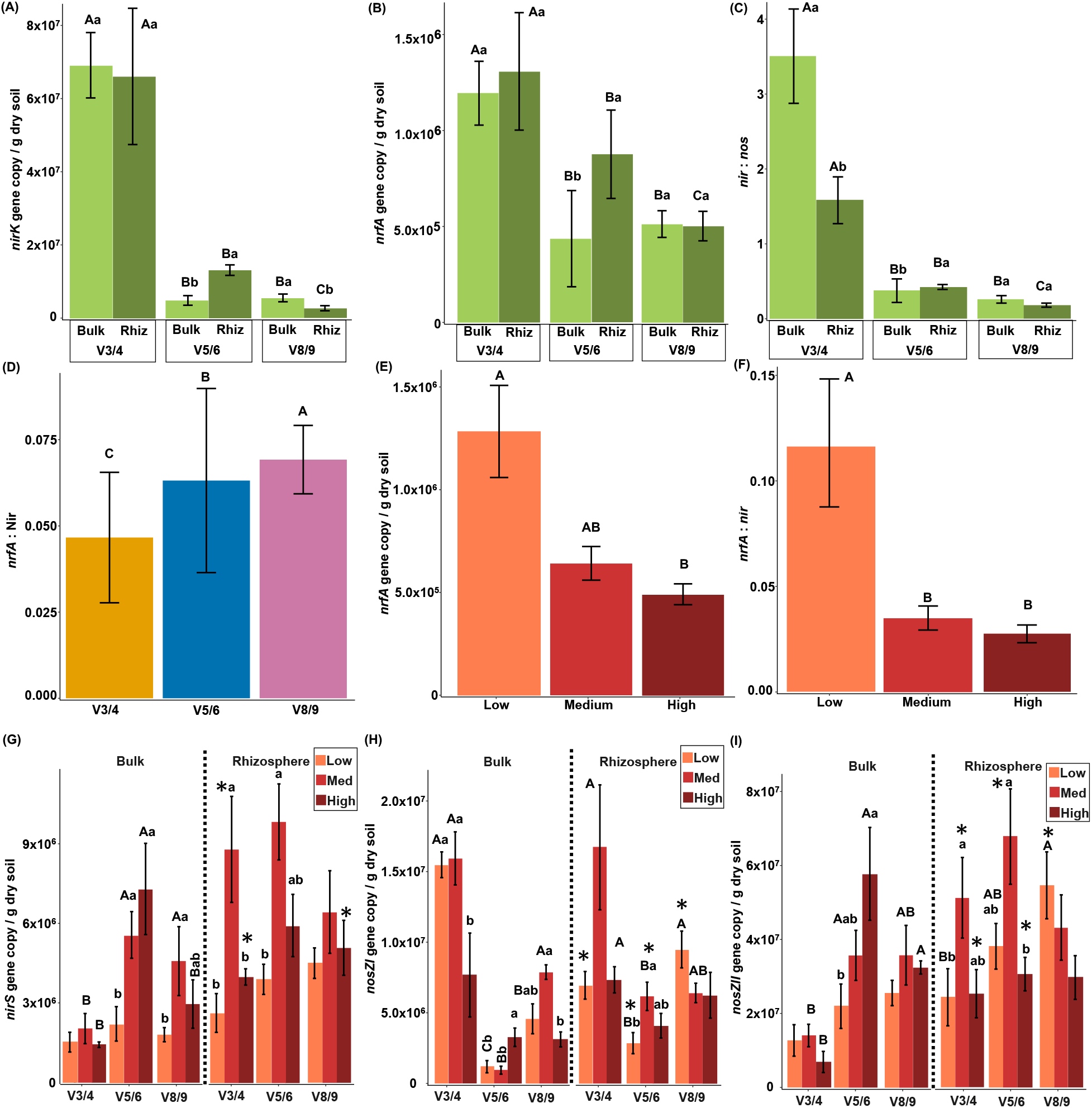
Gene abundances for denitrifiers and nitrate ammonifiers from corn entries. Abundances of nirK (A), nrfA (B), and nir: nos (C) between growth stage x soil compartment and nrfA : nir by growth stage (D). Abundances of nrfA (D) and nrfA : nir (E) by tillage intensity treatments. Abundances of nirS (G), nosZI (H) and nosZII (I) by growth stage x tillage x soil compartment. Uppercase letters in A-F indicate differences between growth stages within soil compartment or tillage treatments and lowercase letters indicate differences by soil compartment within growth stages. Uppercase letters in G-I indicate differences in gene abundances between growth stages within tillage intensity treatments and soil compartment while lowercase letters denote differences by tillage treatments within growth stages and soil compartment and asterisks denote differences in abundances by soil compartment within growth stages and tillage treatments. Error bars represent standard errors.

**Table 4.** Results of linear mixed effect models for N gene abundances from the corn year. Values presented are the F- and p-values for the linear models and significant variables are denoted with an asterisk (*).

| Variable | *nirS* | *nirK* | *nosZI* | *nosZII* | *nrfA* | *nir: nos* | *nrfA : nir* |
| --- | --- | --- | --- | --- | --- | --- | --- |
| Stage | 17.05, <0.01* | 72.82, <0.01* | 44.78, <0.01* | 15.97, <0.01* | 16.77, <0.01* | 79.10, <0.01* | 10.16, <0.01* |
| Tillage | 1.78, <0.01* | 1.34, 0.27 | 3.14, 0.05 | 4.74, 0.01* | 5.56, <0.01* | 3.21, 0.07 | 8.98, <0.01* |
| Compartment | 25.49, <0.01* | 0.08, 0.78 | 11.32, <0.01* | 18.97, <0.01* | 6.72, 0.01* | 0.69, 0.42 | 1.93, 0.18 |
| Stage x Tillage | 1.69, 0.17 | 1.53, 0.21 | 5.10, <0.01* | 1.84, 0.13 | 0.80, 0.53 | 1.84, 0.14 | 0.36, 0.27 |
| Stage x Compartment | 4.08, 0.03* | 13.42, <0.01* | 6.90, <0.01* | 3.40, 0.04* | 7.03, <0.01* | 8.15, <0.01* | 0.74, 0.48 |
| Tillage x Compartment | 0.81, 0.46 | 0.31, 0.73 | 0.43, 0.65 | 4.57, 0.01* | 0.50, 0.61 | 1.46, 0.26 | 0.31, 0.74 |
| Stage x Tillage x Compartment | 3.21, 0.02* | 0.98, 0.43 | 4.84, <0.01* | 2.82, 0.03* | 0.57, 0.69 | 0.39, 0.82 | 0.40, 0.81 |

### Denitrification genes across soybean growth stages

Differences in denitrifier and nitrate ammonifier gene abundances were observed for growth stages, tillage intensities, and soil compartments, but differences were not observed in the ratios of *nir* to *nos* gene markers (Table 5). Abundances of *nirS* and *nos* genes increased over soybean growth stages in both rhizosphere and bulk soils, while *nirK* genes increased with growth stage only in the rhizosphere samples (Figure 5). In bulk soils, the ratio of *nrfA* : *nir* was highest at the last growth stage, R1. Both *nirS* and *nosZI* gene abundances were highest in soybean rhizospheres in the CD tillage treatment. Gene abundances for *nosZI* were higher in soybean rhizospheres than in bulk soils only in the NT treatment (Figure 5).

**Figure 5.**
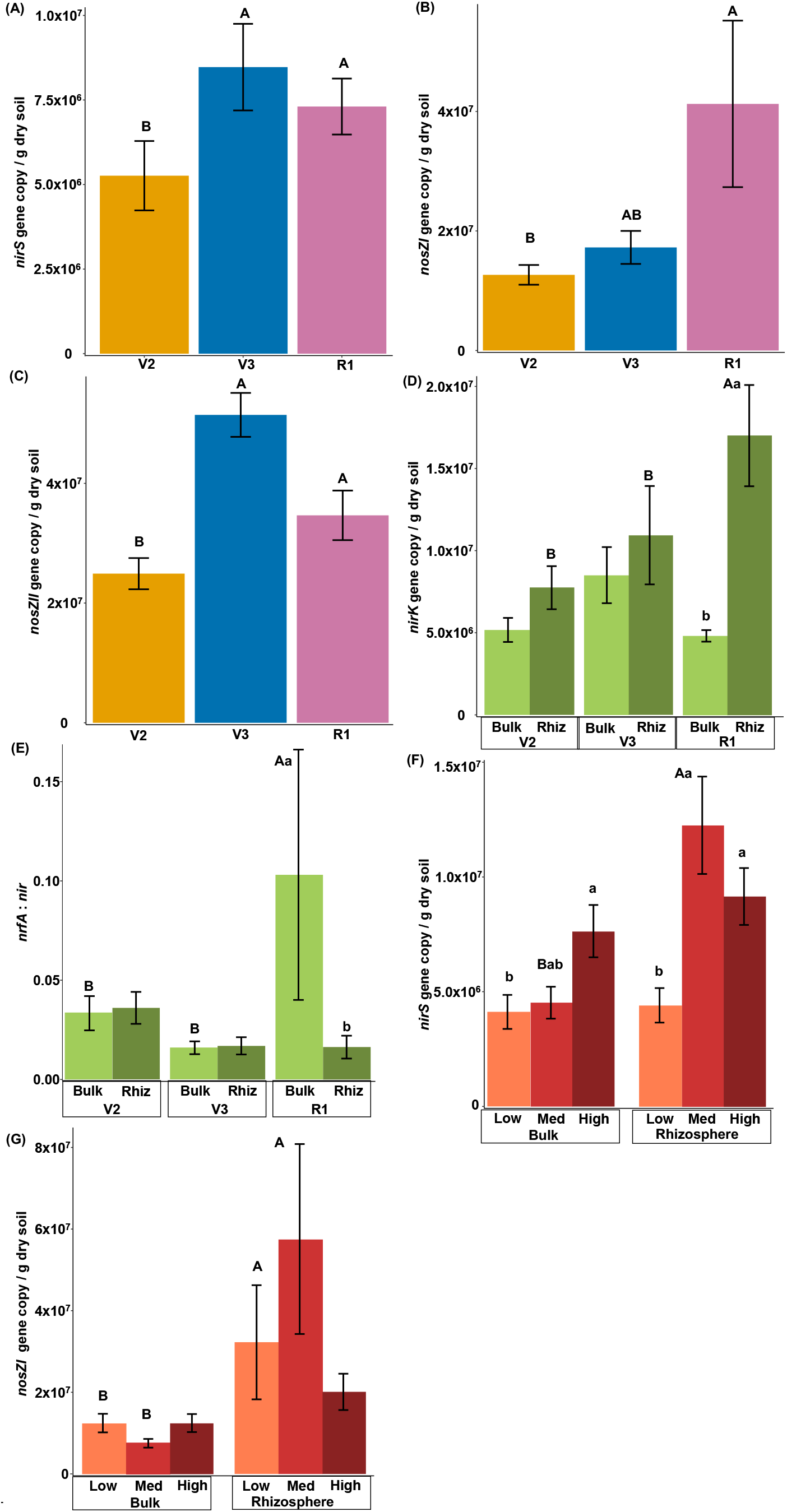
Abundances of genes representing denitrifiers and nitrate ammonifiers from soybean entries. Abundances of nirS (A), nosZI (B), and nosZII (C) over soybean growth stages. Abundances of nirK (D) and nrfA : nir (E) between stage x soil compartment and nirS (F) and nosZI (G) between tillage treatments x soil compartments. Uppercase letters indicate differences in gene abundances between tillage intensity treatments and growth stages and lowercase letters indicate differences between soil compartments in A-E. Uppercase letters in F-G indicate differences in gene abundances by soil compartment grouped within tillage treatments and lowercase letters indicate differences in gene abundances between tillage treatments grouped within soil compartments.

**Table 5.** Results of linear mixed effect models for N gene abundances from the soybean year. Values presented at the F- and p-values for the linear models and significant variables are denoted with an asterisk (*).

| Variable | *nirS* | *nirK* | *nosZI* | *nosZII* | *nrfA* | *nir: nos* | *nrfA : nir* |
| --- | --- | --- | --- | --- | --- | --- | --- |
| Stage | 8.16, <0.01* | 2.10, 0.14 | 5.62, <0.01* | 13.67,<br><0.01* | 2.05, 0.14 | 1.04, 0.36 | 4.96, 0.01* |
| Tillage | 13.08, <0.01* | 0.96, 0.41 | 0.19, 0.82 | 0.01, 0.99 | 0.43, 0.66 | 1.56, 0.23 | 1.42, 0.25 |
| Compartment | 13.16, <0.01* | 9.88, <0.01* | 29.27, <0.01* | 0.76, 0.39 | 0.24, 0.63 | 0.41, 0.52 | 4.39, 0.04* |
| Stage x Tillage | 0.40, 0.80 | 0.77, 0.56 | 0.29, 0.88 | 0.70, 0.60 | 0.72, 0.58 | 1.07, 0.39 | 1.04, 0.40 |
| Stage x<br>Compartment | 1.18, 0.32 | 4.78, 0.02* | 1.65, 0.21 | 1.94, 0.16 | 2.64, 0.09 | 0.14, 0.87 | 5.05, 0.01* |
| Tillage x<br>Compartment | 5.41, <0.01* | 0.29, 0.75 | 3.88, 0.03* | 0.52, 0.60 | 2.19, 0.13 | 0.01, 0.99 | 0.73, 0.49 |
| Stage x Tillage x<br>Compartment | 1.16, 0.34 | 0.14, 0.97 | 1.14, 0.35 | 0.34, 0.85 | 0.68, 0.61 | 0.63, 0.64 | 0.81, 0.53 |

### Management, environmental interactions, and denitrification genes

Conditional inference trees predicting the abundances of each denitrifier gene in response to management and environmental variables are shown in Figure 6. The fixed effects of the experimental design included crop year, tillage intensity, and soil compartment, while environmental variables included precipitation (either one, two, or three days before sampling); mean daily air temperature; growing degree days (GDD); and soil moisture content. At the bottom of each tree, terminal nodes show mean N gene abundances and the number of samples in the group. Based on calculated R^2^-values (Figure 6), the amount of variation in gene abundances explained by the trees was as follows: *nirK* > *nirS* > *nosZII* > *nosZI* > *nrfA*. Management practices including tillage intensity and crop year were the most influential predictors in explaining variation in *nirS* and *nosZI* gene abundances, while climate predictors including growing degree days, air temperature, and precipitation two days before sampling were more influential in explaining variation in *nirK*, *nosZII*, and *nrfA* (Figure 6, node 1). Interactions between management, soil compartment, and/or climate were important in explaining abundances of every gene with the exception of *nrfA*. Abundances of *nosZII*, for example, differed by soil compartment in soybean but in corn abundances varied by air temperature *x* tillage and air temperature *x* growing degree days *x* soil compartment interactions (Figure 6C).

**Figure 6.**
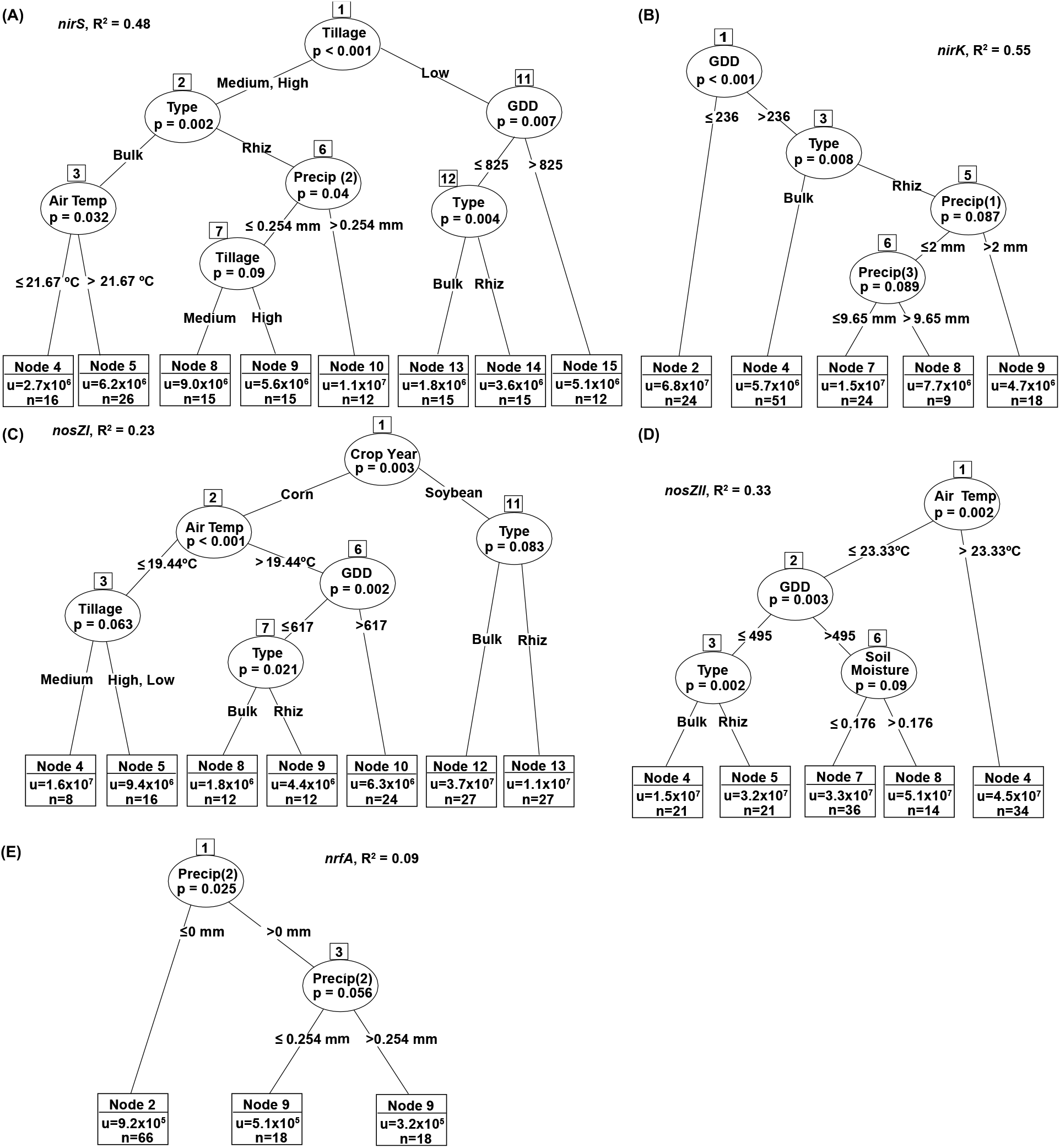
Conditional inference trees explaining the effect of environmental variables (precipitation 1, 2, or 3 days before sampling), mean daily air temperature, growing degree days (GDD; includes corn for 2018 and soybean for 2019), soil moisture content, and fixed effects of the experimental design to include crop year, tillage intensity, and soil compartment (type) to predict nirS (A), nirK (B), nosZI (C), nosZII (D), and nrfA (E) gene abundances. Each node in circles represents the variable that was split, and p-values associated with the split. Values or groups that were split for each variable are shown below each circle. Terminal nodes (in the boxes at the bottom of each tree) includes the mean N gene abundance and number of samples grouped within the splitting criteria. Tree performance was assessed to determine how much variation in the data was explained by the variable splits and are shown as R^2^-values.

## Discussion

The overall objective of this study was to evaluate bacterial-archaeal diversity and composition and denitrification gene abundances in soils subjected to three long-term tillage treatments of increasing disturbance intensity--NT (low), CD (intermediate), and MP (high). Forty years after the experiment’s establishment, we sampled soils from the corn and soybean years of a three-year crop rotation of corn-soybean-small grain + winter cover. Bulk and rhizosphere soils were sampled (depth of 15 cm) at three growth stages of corn and three growth stages of soybean. For 40 years under NT, surface soils would have remained in place and hosted annual root turnover beneath decomposing crop residues. Under CD, soil at the 15-cm depth would have been mixed annually with shallow roots and crop residues, while under MP, soil at this depth would have been replaced and mixed each year with deeper soil and root biomass. We found that bacterial-archaeal assemblage composition and denitrification gene abundances differed between NT and MP treatments, but they did not differ when compared to CD. Community composition and denitrification gene abundances differed between crop types and soil compartment (bulk versus rhizosphere). Bacterial composition and denitrification gene abundances also changed across crop growth stages but were dependent on tillage practices and/or soil compartments.

### Tillage intensity, bacterial-archaeal diversity, and denitrification gene abundance

Tillage explained more of the variation in community composition (R^2^ = 0.07) than either crop or soil compartment, both of which had R^2^ = 0.04. Another long-term tillage study conducted in Illinois, USA, compared bulk soil communities in soybean from NT and conventionally tilled (MP) treatments (Srour et al, 2020). Soils from the MP treatment had higher alpha-diversity and community composition differed between the tillage treatments. In our study, the assessment of both bulk and rhizosphere soils across two crop years (corn and soybean) showed the interactive effects between tillage, soil compartment, and crop type. Tillage intensities appeared to have a stronger influence on assemblage composition than on alpha-diversity across this experiment.

We observed differences in community structure at the family-level based on tillage treatment. Several of these families include taxa that are related to N cycling. Members of the families Nitrosomonadaceae and Nitrospiraceae had higher abundances in NT bulk soils compared to CD or MP soils. Nitrosomonadaceae bacteria oxidize ammonia and are important in the nitrification process (Prosser et al., 2014). Concordantly, the product of ammonia oxidation, nitrite, can be oxidized by members of the Nitrospiraceae family or by some members of Nitrospira genus, which can perform both ammonia and nitrite oxidation to produce nitrate (Daims, 2014; van Kessel et al., 2015). This newly produced nitrate can then be used by organisms performing anammox, the oxidation of ammonium to N_2_ (Kartal et al., 2007). Additionally, Obscuribacteria, a family within the phylum Cyanobacteria; Nitrosotaleaceae a family of archaea that can oxidize ammonia (Prosser and Nicol, 2015); and Phycisphaeraceaea, a family potentially involved in anammox (Rios-Del Toro, et al., 2018), were all higher in NT treatments compared to MP. Increased activity of free-living N_2_-fixing microorganisms may also occur in NT soils (Franzen et al., 2019). Changes to community composition in bulk soils managed under NT favor bacteria and archaea that cycle N and bypass the denitrification pathway. Therefore, managing soils under NT may reduce losses of N as N_2_O or NO_3_^-^ via leaching.

Assessment of N cycling gene abundances further confirmed that NT can increase microbial N use by nitrate ammonifiers and possibly reduce denitrification. Several studies have assessed how microbial N cycling gene abundances are affected when soils are managed with long-term tillage treatments, but none of these studies assessed the effect of tillage on all groups of NO_2_^-^ reducers (*nirS* and *nirK*) and/or both sets of N_2_O reducers (*nosZI* and *nosZII*) (Melero et al., 2011; Tellez-Rio et al., 2015a; Tellez-Rio et al., 2015b; Behnke et al., 2020). Several studies have reported that abundances and activity of *nirS*-denitrifiers and *nirK*-denitrifiers or *nosZI*-N_2_O reducers and *nosZII*-N_2_O reducers have different responses to soil management or environmental variables (Yang et al., 2017; Herold et al., 2018; Hou et al., 2018; Ai et al., 2020; Wang et al., 2021), indicating the need to assess both gene markers to fully understand how management impacts denitrification and N_2_O production/reduction. Our study confirms that these gene sets are not only differentially impacted by tillage intensity but affected in different ways by interactions between tillage, climate, and plant specific rhizosphere selection effects.

### Crop type, community diversity and denitrification genes

We also assessed the effect of crop type on community diversity and N cycling genes associated with denitrification and nitrate ammonification. Several studies have suggested that long-term management practices may have a larger impact and even eclipse the influence of crop species on soil microbiomes (Buckley and Schmidt, 2001, 2003; Jangid et al., 2011). Bulk soils collected from corn and soybean did not show differences in diversity or composition (Chamberlain et al., 2020; Smith et al., 2016), while rhizospheres of soya bean and alfalfa grown in a greenhouse experiment did indicate differences in community composition and alpha-diversity (Xiao et al., 2017). Herein, soybean rhizospheres appeared to have the lowest alpha-diversity and had unique community compositions that were clearly discernible from the soybean bulk soils. Our results indicate that crop species can influence the composition of assemblages, but soil compartment must be considered, as the differences in assemblages were most discernible in the crop rhizospheres.

Rhizosphere, as a habitat, not only affects community composition but also affects gene abundances representing denitrifiers and nitrate ammonifiers. In a greenhouse experiment comparing soil type *x* soil compartment *x* crop type interactions, soil type and soil compartment were the most influential factors affecting N cycling gene abundances (Graf et al., 2016). Plant-microbe interactions in rhizosphere soil is affected by the quantity and type of root exudates which can be unique for each plant species (Berg et al., 2009), water availability, and plant soil N uptake. Therefore, it is likely that microbial N cycling in rhizosphere soils may be constrained by different factors than bulk soils. An example of this is shown in the conditional inference tree for *nirS*, whereby *nirS* abundances in bulk soils are affected by air temperature (Figure 6A, node 2-3) and abundances in the rhizosphere are affected by precipitation and tillage (Figure 6A, node 2, 6, and 7).

### Crop growth stages, on assemblage diversity and denitrification genes

A final aim of this study was to better understand how bacterial-archaeal assemblages and gene abundances for denitrifiers and nitrate ammonifiers change during the growth of corn and soybean. Changes in assemblage composition and alpha-diversity of corn and soybean have been assessed previously (Sugyiyama et al., 2014; Cavaglieri et al., 2009; Xu et al., 2009; Hsiao et al., 2019; Zhang et al., 2012), but not in the context of long-term differences in tillage. Growth stage and tillage had an interactive effect on assemblage diversity in corn, similar to the results observed elsewhere assessing wheat (Wang et al., 2020) and soybean microbiome diversities (Longley et al., 2020). While Longley et al. (2020) analyzed soil microbiomes from different soil compartments, the three-way interaction of soil compartment *x* management *x* growth stage was not presented, so it is difficult to determine exactly how plant selection of rhizosphere microbial assemblages differed from bulk soils over time in their study. Soil compartment and management can synergistically shape rhizosphere communities (Schmidt et al., 2019) and selection strength of rhizosphere microbial communities has been demonstrated to differ between crop species (Tkacz et al., 2015; Berendsen et al., 2012; Uksa et al., 2014). Results from this study indicate that soybean elicits stronger rhizosphere selective effect on bacterial-archaeal assemblages and diversity compared to corn.

By sampling at different growth stages of two crops, we were able to further identify potential ‘hot’ moments of denitrification. Our results support an interaction between management *x* crop rhizosphere selection on N cycling gene abundances proposed by Schmidt et al. (2019). It should be noted that our study and that of Schmidt et al. (2019) both compared management strategies that have been in place for over 20 years. Identifying how management strategies, both in the short-term and long-term, affect locations in the soil (bulk or rhizosphere) and times (growth stages) when denitrifiers are most abundant can provide insights into their changes in activity and further elucidate when and where N_2_O is produced and consumed. This information can be used to target specific soil compartments and growth stages of crops to identify ‘hot spots’ or ‘hot moments’ of N_2_O production and loss.

Finally, results from the conditional inference tree analysis indicate the importance of environmental variables in constraining management and plant rhizosphere selection effects on denitrifier abundances. For example, mean abundances of *nirS* in rhizospheres in the CD and MP tillage treatments were affected by precipitation two days prior to sampling, with rainfall > 0.254 mm resulting in an increase of ∼ 4 x 10^6^ gene copies g^-1^ dry soil. Climate and climate-induced changes to soil conditions have been shown to be highly correlated with denitrification rates in agricultural soils (Kou et al., 2019; Dong et al., 2018; Xu et al., 2020; Saha et al., 2021). Our analyses indicate that climate is not only significantly correlated with denitrification gene abundances but interacts with management practices to influence gene abundances. More intensive tracking of climate and climate-induced changes associated with management variables in modeling analyses should help to provide explanations for high variability in soil N cycling.

## Conclusion

Adoption of reduced tillage practices is a widespread adaptation to global change in agriculture, but it may be associated with environmental tradeoffs of nutrient accumulation and N_2_O emissions. This study demonstrated that management decisions regarding tillage intensity and crop choice have impacts on soil bacterial-archaeal assemblages and denitrification genes. Plant selection strength of rhizosphere assemblages appeared to be stronger in soybean than in corn. Timing of sampling also was an important factor, as community diversity and N gene abundances changed over the crop growth stages. By taking a machine learning approach we were able to identify interactions between tillage practices, growth stages, soil compartments, and climate variables to further demonstrate that gene abundances associated with microbial N cycling were influenced by different combinations of factors. Soils managed with NT had lower gene abundances representing denitrifiers and N_2_O-reducers and greater abundances of taxa associated with other N processes as nitrate ammonification, anammox, and nitrification. Reducing agricultural soil disturbance intensity may provide opportunities to limit N_2_O emissions if we can elucidate the factors that contribute to hot spots or hot moments of N losses.

## Acknowledgements

This research was partially funded by USDA-NIFA grant 1009145 and NESARE grant GNE18-168. Field plots were maintained by the outstanding personnel at the Penn State University Agronomy Farm located at the Russell E. Larson Agricultural Research Farm at Rock Springs. Tyler Bailey was instrumental in helping with field sampling and processing soils in the laboratory.

## Notes

https://github.com/maracashay/Tillage-16SrRNA-MiSeq

### Competing Interest Statement

The authors have declared no competing interest.

