## Supplementary Material for "Tillage intensity and plant rhizosphere selection shape bacterial-archaeal assemblage diversity and nitrogen cycling genes"

Supplementary Material, Tillage management and plant rhizosphere selective effects on bacterial assemblage diversity and nitrogen cycling genes

Supplementary Table 1. Mean average soil fertility results across the three tillage treatments in 2018 (4 blocks sampled) and 2019 (3 blocks sampled). Values presented in the linear model section are the F-values and significant values (p < 0.05) are denoted with an *.

| **Tillage** | **Year** | **pH** | | **Ca (ppm)** | **Cu (ppm)** | **K (ppm)** | **Mg (ppm)** | **P (ppm)** | **S (ppm)** | **Zn (ppm)** | **Acidity** | **CEC** | **K%** | **Mg%** | **Ca%** |
| --- | --- | --- | --- | --- | --- | --- | --- | --- | --- | --- | --- | --- | --- | --- | --- |
| **NT** | **2018** | 6.29 | | 1061.30 | 1.03 | 61.75**a** | 98.25 | 27.50a | 9.28**ab** | 1.20**ab** | 3.10 | 9.38 | 1.68**a** | 8.70 | 56.40 |
| **CD** | **2018** | 6.50 | | 1102.45 | 1.03 | 67.00**ab** | 127.00 | 27.25aǂ | 9.68**a** | 1.25**a** | 2.83 | 9.58 | 1.80**a** | 11.05 | 57.80 |
| **MP** | **2018** | 6.56 | | 1208.85 | 1.08 | 76.00**b** | 148.00 | 22.75aǂ | 8.48**b** | 1.05**b** | 2.05 | 9.50 | 2.03**b** | 12.33 | 62.88 |
| **NT** | **2019** | 6.44 | | 1288.60 | 1.07 | 67.33 | 128.67 | 26.67A | 8.87 | 1.17 | 1.87 | 9.57 | 1.77 | 11.10 | 66.87 |
| **CD** | **2019** | 6.64 | | 1215.10 | 1.13 | 70.67 | 218.67 | 21.67A | 9.00 | 1.27 | 2.63 | 10.73 | 1.63 | 16.57 | 56.53 |
| **MP** | **2019** | 6.96 | | 1385.30 | 1.37 | 117.00 | 293.00 | 15.33B | 7.53 | 0.93 | 1.13 | 10.77 | 2.67 | 21.20 | 63.27 |
| **Linear model** | |  | |  |  |  |  |  |  |  |  |  |  |  |  |
| **Tillage** | | | 1.20 | 0.75 | 1.64 | 4.86* | 1.77 | 23.76* | 4.69* | 5.40* | 1.12 | 1.49 | 7.17* | 1.51 | 0.88 |
| **Year** | | | 1.67 | 3.48 | 3.27 | 4.82* | 4.18 | 34.59* | 7.61* | 0.90 | 1.87 | 5.64* | 3.48 | 3.59 | 1.00 |
| **Tillage x Year** | | | 0.17 | 0.11 | 0.79 | 1.78 | 0.51 | 5.77* | 0.17 | 0.32 | 0.22 | 0.87 | 2.33 | 0.33 | 0.94 |
| Bolded lowercase letters indicate differences between tillage practices averaged across both years | | | | | | | | | | | | | | | |
| ǂ Differences in year averaged across tillage practices | | | | | | | | | | | | | | | |
| Lower case letters indicate differences between tillage practices within the 2018 corn year | | | | | | | | | | | | | | | |
| Upper case letters indicate differences between tillage practices within the 2019 soybean year | | | | | | | | | | | | | | | |

Supplementary Table 2. Primer amounts, qPCR conditions, efficiencies of qPCRs and standards used for qPCR.

| **Gene** | **Primer amount** | **Thermocycling profile** | **Melt Curve** | **Efficiency** | **Standard** |
| --- | --- | --- | --- | --- | --- |
| ***nrfA*** | 1.6 µL of 10 µM (final: 0.8 uM) | 10 min at 95°C followed by 40 cycles of 95°C for 30 s, 51°C for 1 min, and 72°C for 45 s | 95°C for 15 s, 51°C for 1 min, 95°C for 30s and 60°C for 15 s. | 73.23-78.95 | *Escherichia coli K-12* genomic DNA |
| ***nosZI*** | 0.6 µL of 10 µM (final: 0.3 uM) | 10 min at 95°C followed by 40 cycles of 95°C for 30 s, 56°C for 1 min, and 72°C for 45 s. | 95°C for 15 s, 56°C for 1 min, 95°C for 30 s, and 60°C for 15s. | 72.87-84.54 | *Pseudomonas fluorescens* genomic DNA |
| ***nosZII*** | 1.2 uL of 25 µM (final: 1.5 uM) | 10 min at 95°C followed by 40 cycles of 95°C for 15 s, 54°C for 30 s, 72°C for 30 s, and 80°C for 30 s. | 95°C for 15 s, 60°C for 1 min, 95°C for 30 s, and 60°C for 15 s. | 77.84-84.44 | *Gemmatinomas aurantiaca* linearized plasmid |
| ***nirK*** | 0.3 µL of 25 µM (final: 0.375 uM) | 10 min at 95°C followed by 40 cycles of 95°C for 10s and 58°C for 30 s. | 95°C for 15 s, 60°C for 1 min, 95°C for 30 s, and 60°C for 15 s. | 94.69-102.34 | gBlock* from *Ensifer meliloti* 1021 |
| ***nirS*** | 0.8 uL of 10 µM (final: 0.4 uM) | 10 min at 95°C followed by 40 cycles of 95°C for 30 s, 57°C for 30 s, and 72°C for 30 s. | 95°C for 15 s, 60°C for 1 min, 95°C for 30 s, and 60°C for 15 s. | 74.50-83.39 | *Ralstonia eutropha* H16 linearized plasmid |
| **16S** | 0.2 µL of 10 µM (final: 0.1 uM) | 10 min at 95°C followed by 30 cycles of 95°C for 15 s, 60°C for 30 s, and 72°C for 30 s. | 95°C for 15 s, 60°C for 1 min, 95°C for 30 s, and 60°C for 15 s. | 91.60-98.08 | *Flavobacterium columnare* linearized plasmid |
| ****nirK* gBlock** | CCCGCACCTGATCGGCGGCCATGGCGACTATGTCTGGGCTACCGGCAAGTTCCGCAATGCTCCGGACGTCGATCAGGAGACCTGGTTCATACCCGGCGGCACGGCGGGCGCTGCCTTCTACACCTTCGAGCAGCCCGGCATCTATGCCTACGTCAACCATAACCTGATCGAGGCATTCGAGCTT | | | | |

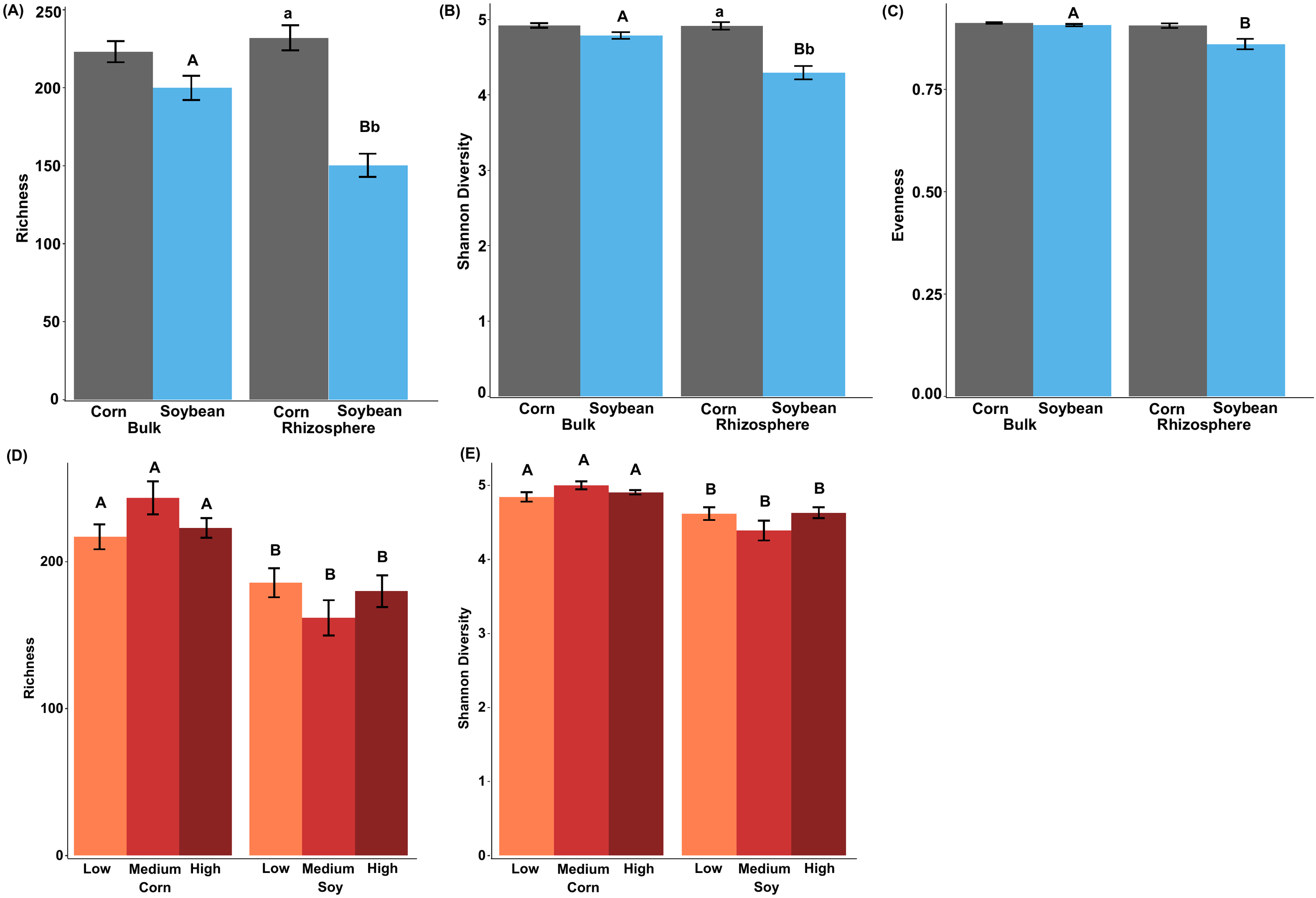
 *Supplementary Figure 1. Alpha -diversity results including richness, Shannon diversity, and evenness. Diversities in the corn and soybean samples analyzed together. Uppercase and lowercase letters denote differences between sample groups. Uppercase letters in A-C indicate differences between soil compartment grouped within crop and lowercase letters indicate differences between crop type (p-values < 0.05). Uppercase letters in D-E indicate differences between crop type.*

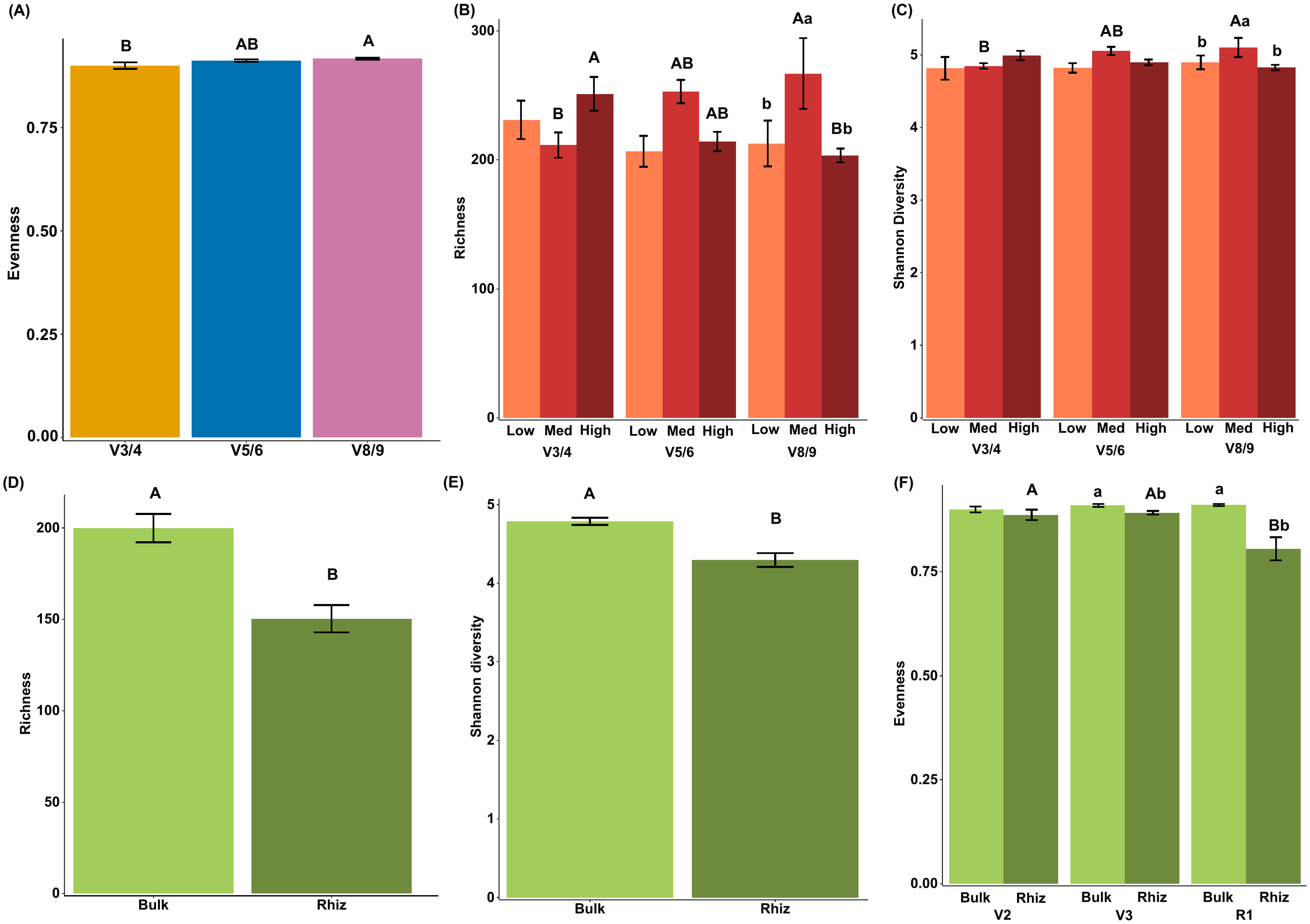

*Supplementary Figure 2. Alpha-diversity results including richness, Shannon diversity, and evenness. Diversity across the corn year to include the three corn growth stages (A-C). Diversities in the soybean year to include the three soybean growth stages (D-F). Uppercase letters and lowercase letters denote which sample groups are significantly different (p-values < 0.05). Uppercase letters in B-C indicate differences across growth stages and lowercase letters indicate differences across tillage practices. Uppercase letters in F indicate differences across growth stages and lowercase letters indicate differences between soil compartments. Error bars represent standard errors.*
